# Asexual male production by ZW recombination in *Artemia parthenogenetica*

**DOI:** 10.1101/2022.04.01.486774

**Authors:** Loreleï Boyer, Roula Jabbour-Zahab, Pauline Joncour, Sylvain Glémin, Christoph R. Haag, Thomas Lenormand

## Abstract

In some asexual species, parthenogenetic females occasionally produce males, which may strongly affect the evolution and maintenance of asexuality if they cross with related sexuals and transmit genes causing asexuality to their offspring (“contagious parthenogenesis”). How these males arise in the first place has remained enigmatic, especially in species with sex chromosomes. Here, we test the hypothesis that rare, asexually produced males of the crustacean *Artemia parthenogenetica* are produced by recombination between the Z and W sex chromosomes during non-clonal parthenogenesis, resulting in ZZ males through loss of heterozygosity at the sex determination locus. We used RAD-sequencing to compare asexual mothers with their male and female offspring. Markers on several sex-chromosome scaffolds indeed lost heterozygosity in all male but no female offspring, suggesting that they correspond to the sex-determining region. Other sex-chromosome scaffolds lost heterozygosity in only a part of the male offspring, consistent with recombination occurring at a variable location. Alternative hypotheses for the production of these males (such as partial or total hemizygosity of the Z) could be excluded. Rare males are thus produced because recombination is not entirely suppressed during parthenogenesis in *A. parthenogenetica*. This finding may contribute to explaining the maintenance of recombination in these asexuals.

## INTRODUCTION

Parthenogenesis usually results in all-female offspring, but many obligate parthenogens are still able to occasionally produce males, such as aphids (Blackman 1972; Rispe et al. 1999; Simon et al. 1999), *Daphnia* (Innes and Hebert 1988), darwinulid ostracods (Smith et al. 2006), nematodes (Snyder et al. 2006), parasitoid wasps (Sandrock and Vorburger 2011), *Potamopyrgus* snails (Neiman et al. 2012) and thrips (van der Kooi and Schwander 2014a). Even though these “rare males” have been reported for a long time, their evolutionary significance remains elusive and controversial (Lynch 1984; Maccari et al. 2013; Engelstädter 2017; Abatzopoulos 2018). If these asexually produced males are non-functional, which apparently is the case in several groups (reviewed in van der Kooi and Schwander 2014b), they can be considered genetic or developmental accidents. The production of non-functional males is costly for parthenogenetic lineages, with the cost depending on the rate at which they are produced (Lynch 1984; Engelstädter 2008; Engelstädter et al. 2011; Neiman et al. 2012). In contrast, if rare males are functional, their evolutionary significance depends on the availability of sexually reproducing females (from related sexual species or lineages, or from their own lineage, if asexuality in females is facultative), with which they can successfully cross and produce fertile offspring. When rare males successfully reproduce with sexual females, they can potentially transmit the gene(s) controlling for asexual reproduction to their offspring, thus generating new parthenogenetic lineages (Hebert and Crease 1983; Simon et al. 2003). Yet whether or not these crosses are successful in generating new asexual lineages and at which rate successful crosses may occur will depend on multiple factors, including the mechanism by which rare males are produced in the first place. Indeed, these mechanisms can be purely accidental or more systematic, the latter generating a regular production of rare males which could be maintained and selected upon. Selection on rare male production could be mediated by selection on the rate of contagion (e.g, in cases where recently derived asexuals enjoy a large fitness advantage). The maintenance of male production may also have important genomic consequences, in particular by contributing to the maintenance of recombination, in cases where the mechanism of male production is based on recombination. Hence, deciphering the mechanism by which males are produced (whether it is accidental or not and whether it is based on recombination) is an important prerequisite to be able to evaluate their possible evolutionary significance.

The possible mechanisms of rare male production strongly depend on the sex determination system occurring in the ancestral sexual species from which parthenogenetic lineages are derived. There are many possibilities, but we opt to classifying them based on genetic patterns that can directly be observed in rare males compared to their parthenogenetic mothers. We distinguish five broad patterns (Fig. 1). Under **Pattern 1**, there is no systematic association between alleles found heterozygous in the mother and the genotype of male offspring. This may occur for instance if males result from accidental phenotypic or genetic “errors” (van der Kooi and Schwander 2014b). For instance, a mutation or transposable element insertion in or near the sex-determining locus may perturb female sex determination of an embryo and result in the production of a male. Similarly, a fortuitous environmental or hormonal variation may induce male development. This occurs for instance with residual environmental sex determination in asexuals derived from species with environmental sex determination (Hebert and Crease 1983; Innes et al. 2000). **Pattern 2** corresponds to the loss of an entire sex chromosome. This may result in male production in species, in which the ancestral sex-determination system is ZZ/ZW, XX/XO, haplodiploidy with a complementary locus (Complementary sex determination, CSD), or depends on X/autosome balance. Mechanistically, the loss of an entire chromosome can occur for instance through non-disjunction. This has been observed in obligately parthenogenetic aphids, where XX mothers produce XO sons (Wilson et al. 1997). Here, it is important to underline that the production of XO male is the “regular” system (Blackman and Hales 1986) in related species of aphids where parthenogenesis alternates with sexual reproduction (cyclical parthenogenesis). **Pattern 3** corresponds to the loss of a part of a sex chromosome (the part that includes the sex-determination locus). This may occur under the same sex determination systems as pattern 2, but the likely mechanism are large-scale deletion or complex chromosomal rearrangement, rather than non-disjunction. This has been observed for instance in stick insects (Pijnacker and Ferwerda 1980). **Pattern 4** corresponds to a complete loss of heterozygosity (LOH) through autozygosity on the sex chromosome pair. Note that we distinguish here LOH leading to autozygosity from hemizygosity (patterns 2 and 3). Pattern 4 may apply to species in which the ancestral sex-determination system is ZZ/ZW, or CSD (Engelstädter 2008). Possible mechanisms include parthenogenesis based on different modifications of meiosis (Archetti 2010; Lenormand et al. 2016). Under these modes of parthenogenesis, all chromosomes (sex chromosomes, as well as autosomes) become homozygous in a single generation. This occurs for instance in rare cases of parthenogenesis in the king cobra, which lead to male offspring, due to terminal fusion (Card et al. 2021). Rare parthenogenesis in heterogametic females leading to the production of males is found in several other species: Komodo dragons where it is associated with complete LOH (Watts et al. 2006), turkey (Olsen and Marsden 1954), and silkworm where induced gamete duplication results in all male offspring (Strunnikov 1995). **Pattern 5** corresponds to partial LOH on the sex chromosome pair. As in pattern 4, it may apply to species in which the ancestral sex-determination system is ZZ/ZW or CSD. Possible mechanisms include modes of parthenogenesis which, in the presence of recombination, lead to partial LOH (Archetti 2010; Svendsen et al. 2015; Lenormand et al. 2016). Under all these mechanisms, the production of rare males occurs due to recombination. For instance, under central fusion or suppression of meiosis I (Archetti 2010), rare males can be produced by LOH at the sex-determination locus through a recombination event between the centromere and this sex-determining locus. This mechanism apparently explains rare male production in CSD species, as in the Cape honeybee (Goudie et al. 2012) and perhaps in *Cataglyphis* ants (Doums et al. 2013). It has also been proposed for ZW *Artemia parthenogenetica* (Browne and Hoopes 1990; Abreu-Grobois and Beardmore 2001; Nougué et al. 2015; Boyer et al. 2021; Rode et al. *in press*) on which we focus in this paper.

**Figure 1.**
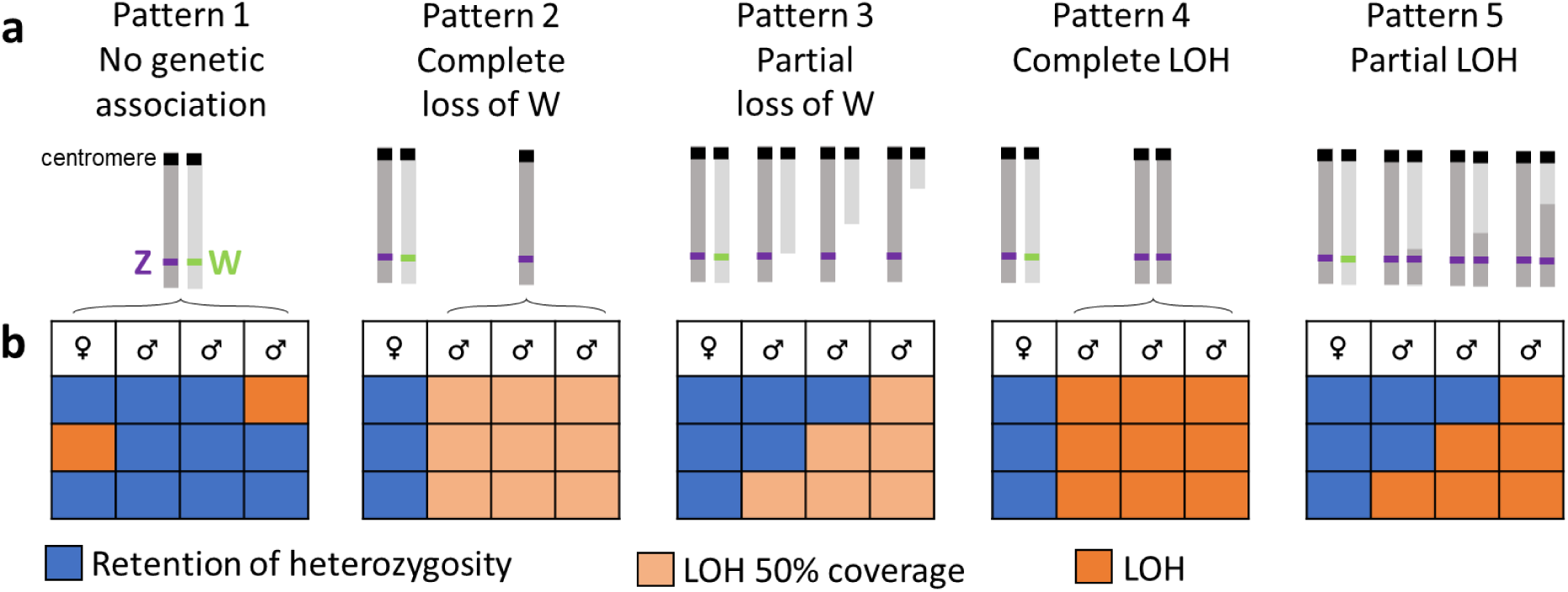
Possible genomic patterns that can be observed in rare males, according to different hypotheses of rare male production. The figure shows the different cases adapted for ZW species, as in *Ap*. (**a**) The ZW chromosome pair is represented with the centromere (black square) and sex-determination locus (purple/green). (**b**) Expectations of LOH patterns on male and female sex chromosome scaffolds (see methods). Each column of the small tables represent individuals (one female and three males). Each line represents a scaffold ordered on the chromosome (top being closer to the centromere). Blue cells represent scaffolds for which maternal heterozygosity was maintained. Dark orange cells represent scaffolds that went through LOH. Light orange cells represent scaffolds that went through LOH and a reduction of 50% of their coverage compared to the scaffolds that retained heterozygosity.

*Artemia parthenogenetica* (hereafter *Ap*) is an obligate asexual species, in which functional males are asexually produced at low rates (Maccari et al. 2013). These rare males are capable of contagious parthenogenesis (Maccari et al. 2014), through mating with females of a closely related sexual species (*A. sp. Kazakhstan*, hereafter *Akaz*, or *A. urmiana*). Moreover, *Ap* females have been shown to rarely engage in sex in the laboratory, which suggests that crosses within asexual populations, through rare males and rare sex in females, might also possible (Boyer et al. 2021). In any case, contagious parthenogenesis was found to contribute to the evolution and diversification of the *Ap* clade (Rode et al. *in press*). Interestingly, the small rate of rare male production seems to consistently differ among *Ap* lineages, perhaps depending on their age (Maccari et al. 2013). Additionally, repeated “contagious” backcrosses of *Ap* to *Akaz* result in an increase in rare male production (Boyer et al. 2021). These observations suggest that rare male production is a genetically controlled trait in *Ap* (MacDonald and Browne 1987; Maccari et al. 2013; Boyer et al. 2021).

In the genus *Artemia*, females are heterogametic ZW, while males are ZZ (Bowen 1963; Stefani 1963). *Ap* are not clonal, but reproduce through a modified meiosis in which the first meiotic division is suppressed but some recombination still occurs (Nougué et al. 2015; Boyer et al. 2021). This mode of parthenogenesis is genetically equivalent to central fusion automixis and leads, as mentioned above, to LOH in the recombinant part of chromosomes, distal (with respect to the centromere) from the position of the crossing-over. Historically, the production of rare male was suggested to occur through the terminal fusion of meiotic products (Stefani 1964), which would lead to complete LOH (pattern 4). Given the more recent findings about the mode of parthenogenesis in *Ap*, rare males may rather be produced by recombination leading to partial LOH (pattern 5) and therefore ZZ offspring at the sex determining locus (Nougué et al. 2015; Boyer et al. 2021; Rode et al. *in press*). However, the sex determination locus and its chromosomal location are unknown, and it is also entirely possible that males are produced by mechanisms not corresponding to this simple scenario. In fact, almost any of the other mechanisms discussed above is possible. Only few can be discarded based on available data. For instance, terminal fusion without recombination is unlikely, as males retain at least some heterozygosity (Abreu-Grobois and Beardmore 2001; Maccari et al. 2013).

In this study, we investigated the mechanism of rare male production by assessing patterns of LOH on the sex chromosomes between mothers and their male and female offspring using RAD-sequencing data. For each of the possible mechanisms for ZW-females, we generated expectations for the patterns 1-5 mentioned above (Fig. 1) for LOH and coverage (to account for possible hemizygosity rather than homozygosity) in rare males, and we then compared observed patterns to these expectations. We also searched for candidate scaffolds for the sex locus in *Artemia* by selecting scaffolds that lost heterozygosity in all male but no female offspring. We discuss the implications of our findings regarding the evolutionary significance of rare males in parthenogenetic species and the maintenance of recombination in parthenogenetic *Artemia*.

## METHODS

### Sex chromosomes in *Artemia*

Generally, the sex chromosomes of *Artemia* are not well characterized. Genetic maps, including Z and W chromosomes, have been obtained in *A. franciscana* (de Vos et al. 2013) and *Akaz* (Haag et al. 2017). Genomic studies have confirmed early cytogenetic observations (Barigozzi 1975), suggesting that the ZW system in *Af* is moderately young and contains a pseudo-autosomal region (de Vos et al. 2013). Two non-recombining parts (Huylmans et al. 2019) show different degree of Z-W divergence. However, the relative size of these regions is unknown. We also do not know whether the sex chromosomes are acrocentric or metacentric (both types are found in *Artemia* chromosomes, Accioly et al., 2014), and the degree of heteromorphy is unclear. Stefani (1963) reported low heteromorphy of sex chromosomes in *A. salina*, W being possibly slightly longer than Z. On the contrary, in *A. franciscana*, high heteromorphy was reported (W being the smallest chromosome, Parraguez et al., 2009), but no heteromorphy was found in another study (Accioly et al. 2014). The position of the sex-determining locus relative to the pseudo-autosomal part of ZW is unknown.

### Sample acquisition and sequencing

The individuals used in this study belong to new parthenogenetic lineages generated by crossing rare *Ap* males with *Akaz* females (Boyer et al. 2021). Since meiosis modification in *Ap* involves suppression of meiosis I with some recombination (Nougué et al. 2015; Boyer et al. 2021), we expect genomic regions far from the centromere to accumulate homozygosity, and therefore to contain no informative markers for LOH in wild lineages (Boyer et al. 2021). Using novel, experimentally generated lineages, rather than individuals sampled from nature, allowed us to circumvent this problem and to investigate LOH in heterozygous markers distributed throughout the genome. Moreover, novel asexual lines generated by hybridization produce rare males at a higher rate than natural *Ap* (6.4 % vs. 0.4 %, Maccari et al. 2013; Boyer et al. 2021). Regarding their sex chromosomes, these new asexual lineages are expected to have a Z-chromosome inherited from *Ap* and a W-chromosome from *Akaz*. Specifically, this study thus investigated whether rare male production in these novel asexual lineages occurs through LOH on sex chromosomes due to recombination between pseudo-autosomal regions of W and Z.

The crosses and rearing conditions have been previously described in detail (Boyer et al. 2021). Shortly, laboratory-maintained *Ap* populations were regularly inspected for rare males. Three rare males were found in a culture of a population sample originally obtained from Aigues-Mortes, France (population PAM7, Nougué et al. 2015) and one from a culture obtained by hatching cysts from lake Urmia, Iran (collection F. Amat, Instituto de Acuicultura de Torre de la Sal, Spain). These four rare males were crossed with five sexual females from *Akaz* (hatched from cycsts, Artemia Reference Center cyst number: ARC1039). The crosses resulted in multiple lineages, which were maintained for several generations of parthenogenetic reproduction. When these lineages produced rare males, we kept the mother, the son, and, when possible, a daughter, for genomic analysis. Across all lineages, nine sons and three of their sisters were used in the analyses, together with all of their mothers (21 individuals in total, fig. S1).

To minimize contamination with non-*Artemia* DNA from microorganisms, individual *Artemia* were washed in two successive baths (10 minutes each) prior to sampling: (1) a 0.05 % sodium hypochlorite solution and (2) sterile salt water (to remove the hypochlorite). Subsequently, the digestive tracks were removed by dissection (to further avoid contamination with gut bacteria), as well as the ovisacs of females (to avoid mixing the DNA of the females with offspring DNA). The resulting tissue was shortly washed in deionized water to remove salt, dried on absorbent paper, and stored in 96% ethanol at −20°C until DNA extraction.

Genomic DNA was extracted using the DNEasy Blood & Tissue kit (Qiagen). Each sample was homogenized in 180 µl of ATL buffer, incubated for 8h at 56°C (700 rpm shaking speed) after addition of 20 µl Proteinase K, and then incubated (5 minutes at room temperature) with 4 µl RNase A. The subsequent extraction steps were carried out according to the manufacturer’s protocol. Library construction and RAD-sequencing were carried out by the Montpellier GenomiX platform (MGX, Montpellier, France). Library construction followed the protocol of Baird et al. (2008), with the restriction enzyme *Pst*I and twelve PCR cycles after ligation of P2 adapters. Sequencing was conducted on an Illumina NovaSeq 6000 flow cell, resulting in 725 million sequences (paired-end, 150 bp).

### Demultiplexing and filtering

We used process-radtags (Stacks v.2.59, Catchen et al. 2013) for demultiplexing, to correct barcodes with one mismatch, and to filter reads with low quality (<0.1 % of reads) or no RAD-tags (11.5 % of reads). We obtained 279 million pairs, with 13 million sequences per individual, on average. Using Trimmomatic v.0.39 (Bolger et al. 2014), we removed cut sites and trimmed all sequences to 139bp. We further retained only reads with quality scores >20 (average) and >15 (all 4 bp windows). This filtering resulted in 10.4% of pairs being removed (because either one or both sequences did not pass), and 250 million pairs being retained.

### SNP calling and VCF filtering

For SNP-calling, we used the de novo/reference hybrid approach as described in Paris et al. (2017) and Rochette and Catchen (2017) in Stacks. We ran “ustacks” on each individual separately, grouping reads that differed by two nucleotides or less into “loci” (parameters: M = 2, m = 3, N = 2). This resulted in an average of 347’564 loci per individual with a mean depth of 23.4 x. We then used “cstacks” to create a catalog of loci across the nine mothers, allowing for a maximum of two differences between individuals for the same locus (parameter: n = 2). This resulted in a catalog of 753’124 loci. The loci of each individual (including offspring) were then compared to the catalog using “sstacks”. On average, 340’601 loci per individual matched with catalog loci. In the next step, we assembled the loci with their paired-end reads using tsv2bam. SNPs and genotypes were called using “gstacks” with default parameters (model: marukilow, parameters: var_alpha = 0.01, gt_alpha = 0.05) and removed PCR duplicates. The latter step resulted in a loss of 78 % of read pairs (PCR or optical duplicates). We retained 36.5 million read pairs, resulting in a total of 733’964 loci with an average depth of 5.7 x, on which genotypes were called.

The catalog loci were mapped to the first-generation draft reference genome of *Akaz*, using bwa mem v.0.7.17 (Li and Durbin 2009), and sorted with samtools v.1.13 (Li et al. 2009). Details of this assembly are given in Appendix 1. We integrated the genomic positions of the loci into the Stacks pipeline using stacks-integrate-alignment with default parameters (mapping quality >20, alignment coverage >60 % and identity percentage >60 %). A large proportion (95 %) of loci mapped to the reference genome, but 32 % of these were removed due to insufficient mapping quality or alignment coverage. The 494’551 remaining loci were filtered using “populations”, keeping only SNPs that were present in more than 13 individuals and removing duplicate loci (separate catalog loci with identical genomic positions). The resulting VCF, containing 359’187 SNPs on 73’402 scaffolds was filtered further, using vcftools v.0.1.17 (Danecek et al. 2011). Given the average depth of 5.7 x, we only kept genotypes based on 2 to 13 reads, thus excluding genotypes with a depth of 1 (which contain no information on whether they are homozygous or heterozygous) as well as genotypes based on more than two times the average depth (because of the elevated risk that these loci included collapsed paralogs). In total, 5.7 million genotypes (on average 271’000 per individual) were retained for the analysis. Fig. S2 shows the depth distribution in the data before filtering.

The sequencing of our library resulted in a high number of duplicate sequences, which were removed as explained above. As a consequence, individual genotype calls were based on a relatively low sequencing depth compared to what is typical for RAD-sequencing data (Rochette and Catchen 2017). Nonetheless, our downstream analysis, investigating LOH between mothers and offspring, shows that even low-coverage RAD-sequencing data can produce highly robust results if the uncertainty of the genotype and the information content of the different loci is adequately taken into account throughout the analysis. Specifically, our analysis is based on genotype likelihoods, rather than fixed genotype calls. These genotype likelihoods were obtained from Stacks. Additionally, we combined likelihood information from all SNPs (minimum = 2) on a given scaffold and used filters to remove data containing little or ambiguous information and to reduce noise. These approaches allowed us to conduct a quantitative analysis of LOH, by propagating uncertainty throughout the analysis rather than overconfidently relying on genotype calls. Similar methods are increasingly used also elsewhere in analysis of genomic data (Korneliussen et al. 2014; Rastas 2017).

### Data analysis: LOH in male and female offspring

The analysis was carried out using the package vcfR v.1.12.0 (Knaus and Grünwald 2017) in R v.4.1.1 (R Core Team 2021) and genotype likelihoods (GL, actually log-likelihoods; Maruki and Lynch 2015; Rochette et al. 2019) obtained as a part of the vcf output in Stacks. Based on the genotype likelihoods (GL), we calculated the probabilities for a given individual (mother or offspring) to be homozygous, *P*_*hom*_, or heterozygous, *P*_*het*_, at a given SNP site. To obtain these probabilities (which have the advantage to sum up to 1), we used exp(GL) to convert the log-likelihoods to likelihoods. Then, we obtained *P*_*het*_ by dividing the likelihood of the heterozygous genotypes by the sum of the likelihoods of all three possible genotypes (because there were two alleles at each site, and thus three genotypes: homozygous for the reference allele, for the alternative allele or heterozygous) and *P*_*hom*_ from 1 – *P*_*het*_.

For each offspring, we restricted our analysis to SNPs for which *P*_*het*_ of the mother was > 0.5. In total, we carried out 766’228 SNP comparisons between mothers and offspring (representing 231’238 SNP localizations on 35’202 scaffolds). For each SNP comparison, we calculated *P*_*LOH*_ (the probability that LOH occurred) by multiplying the *P*_*het*_ of the mother with the *P*_*hom*_ of the offspring. To remove ambiguous SNP comparisons, we removed those with *P*_*LOH*_ between 0.2 and 0.8. For reads containing more than one SNP, we then combined the information by calculating the average *P*_*LOH*_ across all SNPs on the read. For the calculation of the average, each SNP was weighted by the *P*_*het*_ of the mother to give more weight to SNPs that were identified as heterozygous in the mother with higher confidence. To remove instances where the different SNPs on the same read gave conflicting information, we removed comparisons with an average *P*_*LOH*_ between 0.2 and 0.8 as for single ambiguous SNPs. Given that recombination is expected to be rare and that the scaffolds of our draft genome are relatively short (9’638 bp, on average), we then proceeded in the same way, combining the information of all loci (individual SNPs or combined SNPs per read) within scaffolds and retaining only scaffolds with at least two loci. The resulting average *P*_*LOH*_ per scaffold was again weighted by the per-locus average *P*_*het*_ of the mother, and ambiguous scaffolds (i.e., those with average *P*_*LOH*_ between 0.2 and 0.8) were removed. In total, we obtained 125’663 per-scaffold average *P*_*LOH*_ estimates, representing 25’032 different scaffolds, whose combined lengths represent 53% of the total assembly length.

### Identifying autosomal and ZW scaffolds

We used three approaches to identify scaffolds on autosomes vs. sex chromosomes. Together, they resulted in the identification of 96 scaffolds on the sex chromosomes and 1’998 autosomal scaffolds (after the removal of 17 scaffolds with conflicting information, i.e., being assigned to sex chromosomes by one method and to autosomes by another). Given a haploid chromosome number of 21 (Barigozzi 1974), this gives an average of about 100 scaffolds identified per autosome. Fig. S3 shows the number of scaffolds identified for Z and autosomes by the different methods.

First, we used a re-analysis of the raw data from the *Akaz* genetic map (Haag et al. 2017), to integrate the map with the genome assembly. We generated a vcf, using the *de novo*/reference hybrid approach in Stacks described above and filtered loci and individuals as in Haag et al. (2017). We established a preliminary correspondence between our markers and those of the Haag et al. (2017) map, based on segregation patterns among the offspring and used R/qtl (Broman et al. 2003) functions ripple and dropone to further order markers along the linkage groups and to remove markers whose segregation patterns did not fit the linkage group. We then combined markers within scaffolds, removing scaffolds with inconsistent markers (i.e., with different markers mapping do different linkage groups), as well as scaffolds mapping to different linkage groups between male and female maps. In total, this procedure resulted in the identification of 22 Z-linked and 360 autosomal scaffolds.

Second, we assessed orthology between transcripts identified by RNA-sequencing of four *Akaz* males and four *Akaz* females with Z-linked genes and autosomal genes in *A. franciscana* (Huylmans et al. 2019). For RNA-sequencing, live *Akaz* individuals were washed for 2 minutes in sterile, deionized water and shortly dried on absorbent paper before they were flash-frozen in liquid nitrogen and stored at −80°C. RNA extraction was performed using the “NucleoSpin RNA Set for NucleoZOL” kit (Macherey-Nagel), following the manufacturer’s instructions. RNA-sequencing was performed by Genewiz (Leipzig, Germany). The RNAseq reads were mapped to the *Akaz* reference genome (with repeat regions masked using RepeatMasker, Smit et al. 2013) using hisat2 (Kim et al. 2019). Protein coding sequences were identified with Augustus (Stanke et al. 2006), and their orthology to genes on *A. franciscana* scaffolds identified as autosomal or Z-linked (Huylmans et al. 2019) was assessed with OrthoFinder (Emms and Kelly 2015). In cases where groups of orthologs were identified (rather than one-to-one orthologs), *Akaz* and *Af* sequences were aligned using MACSE (Ranwez et al. 2011), and synonymous distance dS was estimated using codeML in PAML (Yang 1997). Only ortholog pairs with the lowest dS were retained for further analysis. In total these procedures allowed identification of 21 Z-linked and 1’670 autosomal *Akaz* scaffolds. *A. franciscana* has an estimated divergence of 32 MY from *Ap* and *Akaz* (Baxevanis et al. 2006). In this analysis, we made the assumption of synteny between *A. franciscana* and *Akaz*, based on the fact that only a single chromosomal rearrangement is known in *Artemia* (Barigozzi 1974). However, to assess the robustness of our results to this assumption, we performed the downstream analyses both including and excluding scaffolds that were identified as sex-linked by this method only.

Third, we used the RNA-sequencing data (see previous paragraph) to identify putative sex-linked SNPs that were consistently homozygous in *Akaz* males (putatively ZZ) and consistently heterozygous (putatively ZW) in *Akaz* females. Reads were mapped to the *Akaz* reference genome with the STAR aligner (Dobin and Gingeras 2015) and duplicates were removed with Picard RemoveDuplicates (Broad Institute). Variants were called with GATK HaplotypeCaller (van der Auwera and O’Connor 2020). Only variants genotyped in all individuals and with a minimal coverage of 10 were retained for this analysis. We selected loci with a minor allele frequency of less than 0.1 (homozygous-tendency) in all males and of more than 0.1 (heterozygous-tendency) in all females. We then identified scaffolds where the frequency of such loci was higher than expected by chance. To do so, we first computed the probability *p*_*zw*_, that a SNP could show by chance a ZW pattern (meaning being homozygous in all males and heterozygous in all females) given the average heterozygosity in males and females (across 200,635 identified SNP with a minimum of 10x depth in every individual). For each scaffold, we observed the number of SNPs showing a ZW pattern and we then tested whether their proportion exceeded *p*_*zw*_ (using a simple binomial test). Finally, we applied a Bonferroni correction across scaffolds for multiple testing. We used this correction for its high stringency. This method allowed us to identify 58 sex-linked scaffolds.

### LOH on sex chromosomes and autosomes

After filtering (see above), we retained a *P*_*LOH*_ estimate for 77 of the 96 sex-linked scaffolds in at least one mother-offspring pair (521 mother-offspring comparisons in total). Considering *P*_*LOH*_ > 0.8 as LOH and *P*_*LOH*_ < 0.2 as heterozygosity retention (i.e., no LOH), we constructed a map of the scaffolds along the sex chromosome by ordering scaffolds according to LOH frequency across mother-offspring pairs. We further arranged the map manually by grouping scaffolds with LOH. To distinguish whether LOH events were due to homozygosity or hemizygosity (see Fig. 1), we compared the depth of the scaffolds that lost heterozygosity with the depth of the scaffolds that retained maternal heterozygosity. To compute this, we first scaled each scaffold depth by the average individual scaffold depth (to correct for small variation of sequencing effort among individuals). Then, we computed the mean scaled depth for ZW scaffold that lost and did not lose heterozygosity.

We investigated whether males and females were produced by meiosis showing different recombination patterns on autosomes. To do so, we compared autosomal LOH rates for male and female offspring. Among the 1’998 autosomal scaffolds, 1’393 were retained for the analysis (9’072 *P*_*LOH*_ estimates for individual mother-offspring comparisons in total). To analyse the data, we fitted likelihood models as described in Appendix 2. Because patterns of autosomal LOH seemed variable among individuals, we fitted a model allowing for inter-individual variation in rates of autosomal LOH. Specifically, we assumed that individual rates of autosomal LOH followed a Beta distribution. Because autosomal LOH rate was particularly variable in males, we then tested whether these distributions differed between males and females using likelihood ratio tests.

### Candidate scaffolds linked to the sex-determining locus

To assess which candidate scaffolds are likely most closely linked to the sex-determining locus, we identified scaffolds for which all male offspring (for which data were available) lost heterozygosity (*P*_*LOH*_ > 0.8) while all female offspring retained heterozygosity (*P*_*LOH*_ < 0.2). Here, we used all scaffolds, regardless of whether we were able to assign them to the sex chromosomes, autosomes, or not. After filtering, a total of 15’337 scaffolds had a *P*_*LOH*_ estimate for at least one male and one female offspring. But among those that fitted the expected pattern (LOH in all male, no LOH female offspring), data were available for only 38% of offspring, on average. We therefore also added cases where the mother genotype was missing or of too low quality, as long as at least one offspring was clearly heterozygous, as this unambiguously indicates heterozygosity retention (and therefore indicates that the mother was heterozygous as well). We used the same quality filters as above (*P*_*LOH*_ lower than 0.2), and obtained 5’631 additional genotypes on 202 scaffolds, which were used to check whether these scaffolds still matched the expected patterns. Two scaffolds were removed from the final candidate list because they were assigned, by above-mentioned approaches, to autosomes.

## RESULTS

### LOH on sex chromosomes

#### Patterns of LOH in male and female offspring

The expected patterns of LOH on sex chromosome scaffolds according to the different hypotheses are represented in Fig. 1b. None of the female offspring lost heterozygosity on any of the 77 sex-chromosome scaffolds. In contrast, all male offspring lost heterozygosity in some of these scaffolds (Fig. 2). LOH on 18 scaffolds (23 % of all scaffolds) was shared among all males, that is all genotyped males lost heterozygosity on these scaffolds, whereas no LOH in any of the genotyped males was observed on 20 scaffolds (26%). For the remaining scaffolds, LOH was variable among males (Fig. 2). With a single exception (one scaffold in male B3 for which we cannot exclude genotyping error or erroneous sex chromosome assignment), the LOH patterns of all scaffolds were consistent with a single crossover having occurred at a variable location between W and Z during the production of every male offspring. Crossovers results in retention of heterozygosity for all scaffolds between the centromere (putative location: top of Fig. 2) and the crossover location, while leading to LOH for all scaffolds between the crossover location and the telomere (putative location: bottom of Fig. 2). The scaffolds in Fig. 2 are therefore likely ordered approximately as they are on the sex chromosomes. The scaffolds for which LOH was consistently observed in all males contain the inferred location of the sex-determining locus (putatively becoming ZZ in all males while remaining WZ in all female offspring). The ratio of the mean scaled depth of scaffolds that lost heterozygosity relative to the depth of scaffolds that maintained heterozygosity was 1.12 ± 0.03, so not consistent with hemizygosity (expected ratio = 1/2). Excluding scaffolds assigned by orthology with *Af* (i.e., only including those identified by the genetic map and the RNAseq studies) reduced the data set to 68 ZW scaffolds but had no qualitative nor major quantitative effect on the results (Fig. S4).

**Figure 2.**
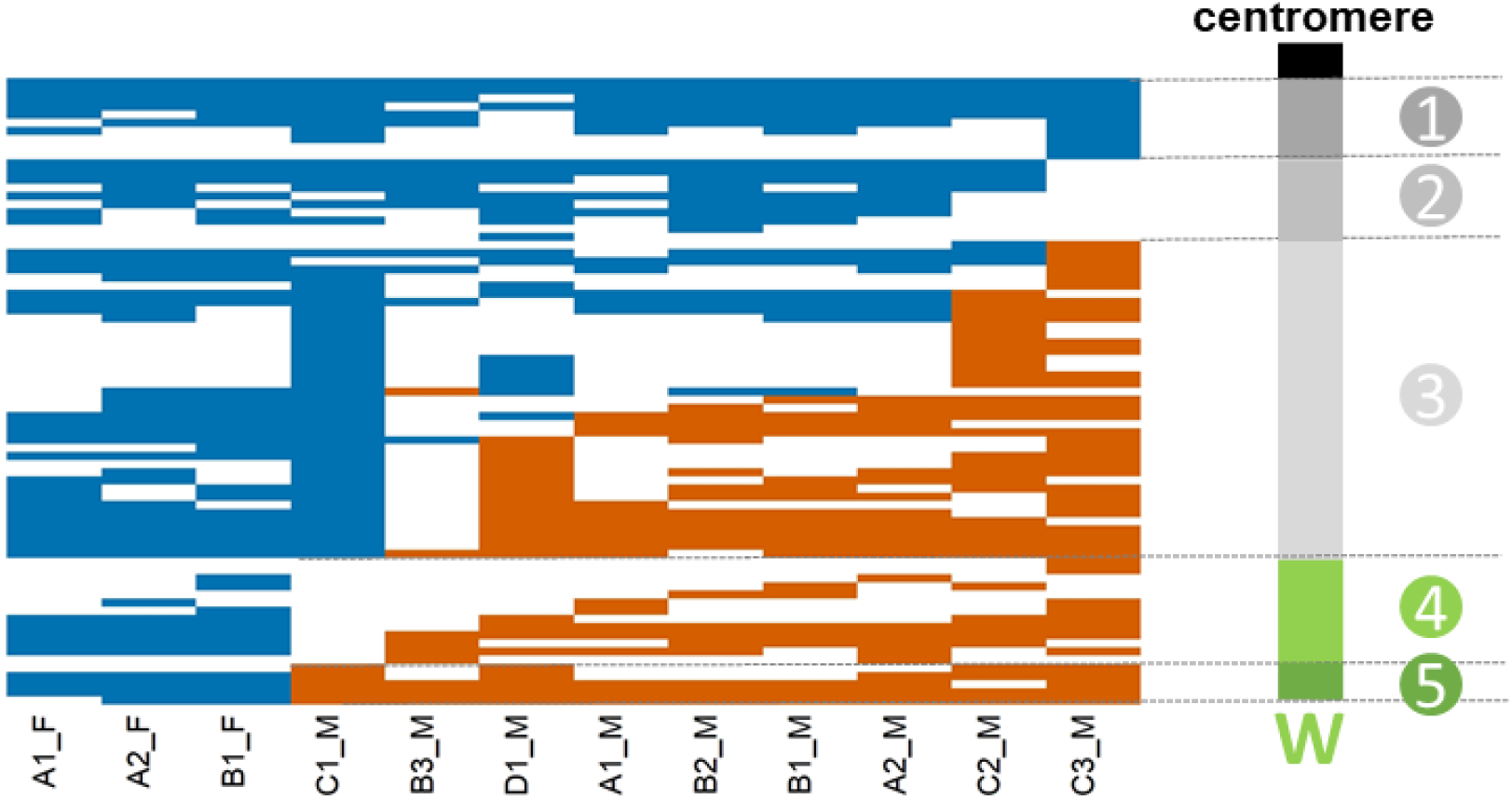
LOH on the 77 sex chromosome scaffolds. Each column represents an offspring, first the three female offspring then the nine male offspring, and each line represents a scaffold. Blue indicates heterozygosity retention (*P*_*LOH*_ < 0.2), orange indicates LOH (*P*_*LOH*_ > 0.8), while white indicates that information is not available for a given scaffold in a given offspring. Scaffolds are ordered from top to bottom according to increasing *P*_*LOH*_ frequency across offspring, and offspring are ordered from left to right according to increasing *P*_*LOH*_ frequency across scaffolds. On the right, a possible reconstruction of the W according to the results. (1) Non recombining region near the centromere (13% of the scaffolds / 17.1% of bp sequenced). This region can be extended to region (2), although information is missing for some scaffolds where recombination could have taken place (the whole region would then be 26% of the scaffolds / 30.2% of bp sequenced). (3) Pseudo-autosomal region where some recombination is observed (51% of the scaffolds / 49.1% of bp sequenced. (5) Non-recombining region around the sex-determination locus (6% of the scaffolds / 8.0% of bp sequenced). This region can be extended to region (4), although information is missing for some scaffolds where recombination could have taken place (the whole region would then be 23% of the scaffolds / 20.7% of bp sequenced).

#### Candidate scaffolds likely linked to the sex-determining locus

When considering all scaffolds with LOH data, that is, including those that could not be assigned to either sex chromosomes or autosomes, and adding the information of scaffolds that were not genotyped in the mother, we identified 58 scaffolds potentially linked to the sex-determining locus. However, two were assigned to the autosomes and were removed. Three of the remaining scaffolds were assigned to the sex chromosomes, and we could not infer the assignation of the others. Given the low number of offspring, these may contain a number of false positives, potentially even including autosomal scaffolds. Nonetheless, with the additional data considered (see methods), an LOH estimate was available for almost 80 % of offspring, on average for a given scaffold.

### LOH on autosomes

The distribution of autosomal LOH among offspring differed between males and females (likelihood ratio test, *P* = 0.002). Autosomal LOH was significantly larger (and more variable) in males than in females. More specifically, in males, LOH rate followed a beta distribution with an estimated mean of 0.07 and variance of 0.01 (a = 0.67; b = 8.41). In females, LOH rate followed a beta distribution with an estimated mean of 4.10^−3^ and a near-zero variance of 10^−9^ (a = 17.6; b = 4.0 ×10^6^; Fig. 3). Excluding scaffolds assigned by orthology with *A. franciscana* (i.e., only including those identified by the genetic map) reduced the data set to 260 autosomal scaffolds but had no qualitative nor major quantitative effect on the results (Fig. S5).

**Figure 3.**
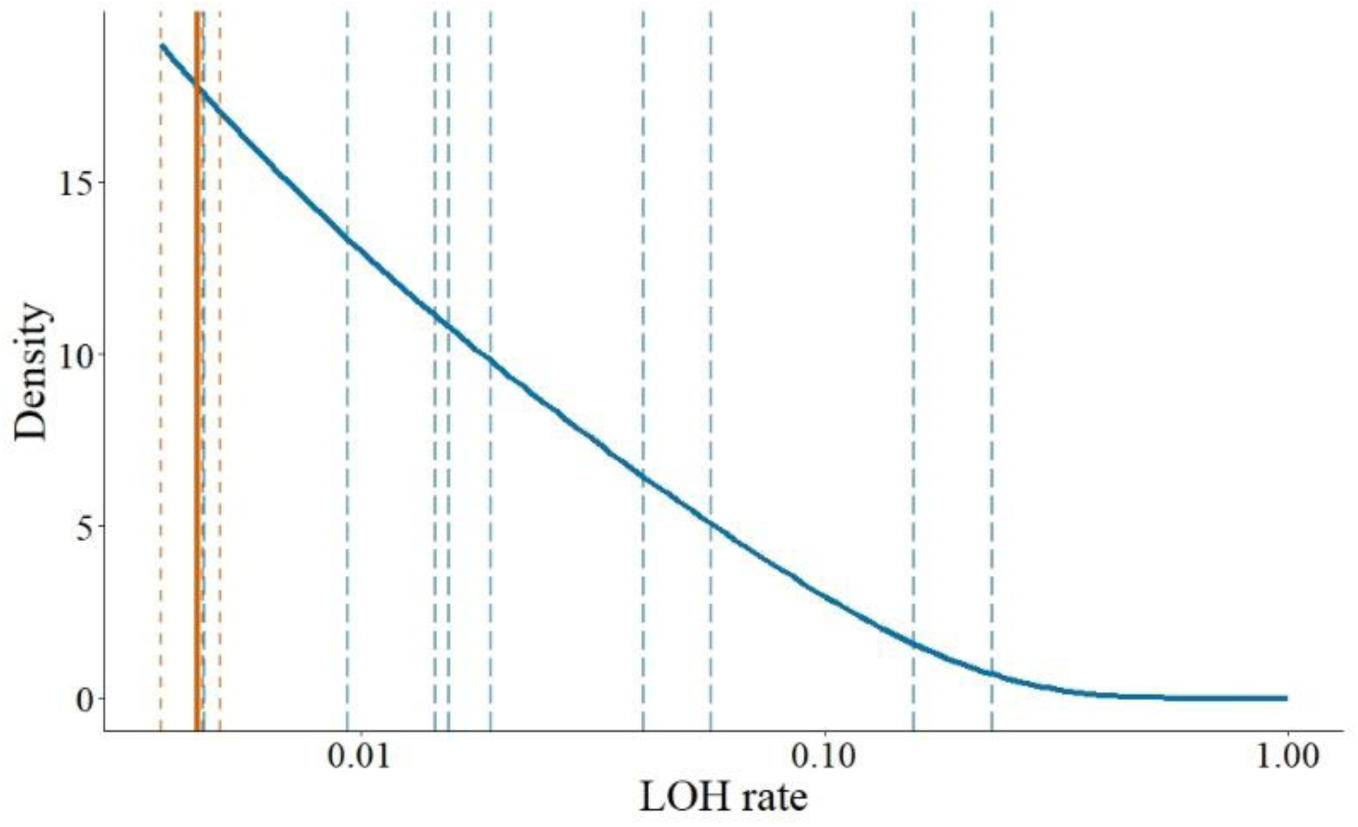
LOH rate on autosomes in male (blue) and female (orange) offspring. Dashed lines represent LOH rates in autosomal scaffolds for individual offspring, and solid lines represent the estimated distributions from our best model. Note the log-scale of the x-axis.

## DISCUSSION

### Rare males are produced asexually by recombination on the sex chromosomes

We found LOH patterns on sex chromosome scaffolds exactly as expected under the hypothesis that rare males result from LOH events due to recombination breakpoints occurring at a variable location between the centromere of the sex chromosome pair and the sex-determining region (pattern 5 in Fig. 1). We found a systematic genetic signature of sex chromosome LOH in all males and absence thereof in all females. This observation excludes environmental or other errors in sex differentiation as well as localized mutation or TE insertion near or at the sex-determination locus as the main explanation for the production of rare males. LOH in males concerned only a part of the sex chromosome scaffolds, while others retained heterozygosity in all males. This observation rules out non-disjunction that would lead to a complete loss of the W (i.e., ZO males) as well as modified meiosis mechanisms (e.g., terminal fusion without recombination) resulting in homozygosity over the entire chromosome. Note that terminal fusion leads to systematic LOH at all scaffolds (no recombination) or at the centromere-proximal scaffolds (with recombination) on all autosomes as well. This mechanism (suggested by Stefani, 1964, for *Ap*) can therefore be excluded according to our results (no systematic LOH on autosomes, neither in males nor in females). Finally, sequencing depth of sex chromosome scaffolds that lost heterozygosity was similar (even slightly higher) than the depth of scaffolds that retained heterozygosity. This strongly suggests that LOH is due to homozygosity rather than hemizygosity and therefore caused by recombination during modified meiosis rather than by partial W deletion.

Given these findings, we can conclude with high confidence that the nine males in our analysis were produced by LOH at the sex-determining locus, caused by ZW crossover events that occurred during modified meiosis in their mother. This mechanism is in line with our knowledge of the asexual reproductive mode of *Ap* (Nougué et al. 2015; Boyer et al. 2021), and has been previously suggested as a possible mechanism for rare male production (Browne and Hoopes 1990; Nougué et al. 2015; Boyer et al. 2021). Recombination on sex chromosomes can lead to ZZ or WW offspring, not for the entire chromosome, but at least for the region relevant for sex-determination. While WW individuals are probably non-viable, ZZ individuals develop as rare males.

Rare males were therefore produced by recombination, but interestingly, it also appears that they were often produced by meiosis showing particularly high recombination rates overall, i.e. even on autosomes (compared to the meiosis leading to female offspring). In autosomal scaffolds, the distribution of LOH rate differed between male and female offspring. Females had a consistently very low rate of LOH while males displayed very variable LOH (up to 16%). Males and females might be the result of reproductive events with different recombination rates. However, since there is a high variation in male offspring LOH, it seems more likely that this difference is actually the result of continuous intra-mother variation in recombination rate, and that males simply happen to be produced by (modified) meiosis having a higher than average number of crossing overs on all chromosomes, including the sex chromosomes. The difference observed between male offspring with the most LOH and female offspring could also be partly due to population or cross variation: Although the females are controls as they are the sisters of males in our analysis, we do not have data for the sisters of the males that happen to show the highest LOH.

While *Ap* females reproduce mainly through modified meiosis, they can rarely reproduce sexually, likely through a normal meiosis (Boyer et al. 2021). In contrast, rare males produced by these *Ap* females seem to mainly or always undergo normal meiosis as no evidence for unreduced sperm (i.e., no evidence for triploidy in their offspring) was found (Boyer et al. 2021). This suggests that both normal and modified meiosis pathways exist in *Ap* and that the frequency of the expression of one or the other pathway is sex-specific.

Finally, our proof that rare males are produced by recombination validates the use of the rate of rare male production as a good proxy to measure residual recombination rate in asexual lineages (Nougué et al. 2015; Boyer et al. 2021). Of course, this is only a measure of the recombination rate on the sex chromosome pair, and it may not necessarily reflect global recombination rate in the genome. However, it may be a less biased estimate for wild populations, as the history of past LOH precludes to estimate LOH rates in an unbiased manner using genetic markers (since the loci away from the centromere, which are the most likely to undergo LOH are unlikely to be heterozygous in the first place).

### Explaining the heterogeneity of rare male production among lineages

Different *Ap* lineages produce rare males at different rates (Maccari et al. 2013). According to our results, this can be explained by these lineages having different recombination rates. Under modified meiosis where meiosis I is suppressed, recombination leads to LOH. Similarly to inbreeding, LOH leads to lower fitness, as it reveals recessive deleterious mutations (loss-of-complementation, Archetti 2004). Hence, it is likely that there is selection for a lower recombination rate within asexual lineages (Engelstädter 2017). This was suggested in the Cape honey-bee, which also reproduce by a modified meiosis where recombination causes LOH (Goudie et al. 2012). Reduction of the recombination rate would result in a lower rate of male production. In *Ap*, this scenario is consistent with the observation that rare male production increases in asexual females obtained from successive backcrosses to the sexual species *Akaz* (therefore introgressing sexual recombination determinants, Boyer et al. 2021). Now that there is conclusive evidence for rare male production by recombination, we can firmly interpret this earlier result as evidence for selection against recombination in asexual lineages compared to the sexual species. In some lineages, in which seemingly no rare males are produced (Maccari et al. 2013), recombination might have been lost altogether (or equivalently the position of crossing-over might have evolved to be telomeric, and thus not cause LOH). Similarly, in *Ap* polyploids, no male has ever been observed. To explain this observation, it is often assumed that these polyploids reproduce by clonal apomixis. This is however unlikely, as they are derived from crosses involving asexual diploids (Rode et al. *in press*). They are therefore likely to share the same meiosis modification as the diploids. The absence of rare males in these lineages may rather result from the preferential pairing of Z with Z and W with W, drastically limiting the opportunity for recombination between Z and W. Furthermore, in tetraploids and pentaploids, two subsequent LOH would be required to produce ZZZZ or ZZZZZ males, which further reduces the likelihood of occurrence of rare males (Rode et al. *in press*).

### Long term evolution of recombination in *Ap* lineages

If LOH is costly, why then is non-zero recombination and with it rare male production maintained in most *Ap* lineages? One possibility is that most lineages are relatively recent, and recent lineages maintain some recombination. This could be because they were recently generated by a sexual event involving a rare male produced by recombination or because of introgression of recombination alleles if the new lineage was created by crossing with a female from a sexual species (Boyer et al. 2021). Consistent with this idea is the observation that the highest rates of rare male production occur in *Ap* lineages from Central Asia (Maccari et al. 2013), that is, from the distribution range of the sexual species, where repeated crosses are most likely to occur. Yet, phylogenetic evidence suggests that crosses with sexual species leading to contagious asexuality occurred only rarely in the evolutionary history of extant diploid *Ap* (Rode et al. *in press*). The generation of new asexual lineages may thus more frequently involve rare males and rare sexual events in *Ap* females species (Boyer et al. 2021). Such within-*Ap* crosses would be more difficult to detect with phylogenetic evidence. Another possibility is that rare within-*Ap* crosses or contagious parthenogenesis might sometimes lead to new asexual lineages with fewer or masked deleterious mutations, thus “rescuing” old asexual lineages from long-term decrease in fitness. However, whether the possibility to generate new asexual lineages via rare males indeed can lead to selection for maintaining non-zero recombination requires further study. It should notably involve a study of the costs of LOH through unmasking recessive deleterious mutations and the possibility that selection pressures may differ for the asexuality-determining region(s) vs. the rest of the genome.

### Structure of the Z chromosomes

The fact that males are produced by recombination means that portions of the W and Z chromosomes are indeed pseudo-autosomal, and that this pseudo-autosomal region (PAR) is located between the centromere and the sex-determining locus. Moreover, the observed heterogeneity of male LOH on the sex chromosomes suggests that crossovers may occur at many different locations and thus that the PAR is relatively large (51-81 % of the sex chromosome scaffolds underwent LOH in at least one male while retaining heterozygosity in at least one other male, suggesting that all these scaffolds are located in the PAR). The non-recombining region near the centromere contains 13%-26 % of all sex chromosome scaffolds with *P*_*LOH*_ data and the non-recombining region containing the sex-determining locus 6% - 23 % of all sex chromosome scaffolds with *P*_*LOH*_ data. Note that it is possible that a part of the PAR is on a second chromosome arm or located terminally after the sex-determining region. However, if present, these regions are likely small (or have a short genetic distance) as otherwise we should have observed sex chromosome LOH in females as well. Fig. 2 represents these findings and a possible structure of the W.

We identified 56 scaffolds potentially associated with the sex-determining locus, but of these, only three scaffolds could be assigned to the sex chromosome pair. Better genomic resources for *Ap* might allow narrowing this list. Nonetheless, these scaffolds constitute an important first step towards identifying the sex-determining locus in *Artemia*.

## Conclusion

In this study, we compared RAD-sequencing data between asexual *Artemia* females and their male and female offspring. We demonstrated that rare ZZ males are produced by recombination between W and Z sex chromosomes, as a result of a non-clonal asexual reproductive mode. The data also allowed us to infer the likely structure of the sex chromosomes, the localization of the sex-determining locus, and a list of candidate scaffolds associated with the sex-determining locus. This study shows that the consequences of non-clonal asexuality, even occurring through rare events, can be significant. Rare males potentially are major actors in the long-term evolution of *Artemia*. Knowing how rare males are produced in parthenogens, when combined with reliable genomics resources, can provide essential insight into their evolutionary significance, and the consequences of contagious parthenogenesis.

## ACKNOWLEDGMENTS

The authors are grateful to G. Van Stappen for providing the *Akaz* sample. We thank E. Ortega, Q. Rougemont and E. Beyne for helpful advice with the bioinformatics analysis. We thank M.-P. Dubois and The Genomics, Molecular Ecology, and Experimental Evolution platform (GEMEX) at CEFE. RAD-sequencing library construction and sequencing was performed by the MGX platform (Montpellier, France). RNA-sequencing was conducted by the Genewiz company, (Leipzig, Germany). We are thankful to the GenOuest bioinformatics core facility for providing computing infrastructure for part of this study. This work was funded by the Grant ANR-17-CE02-0016-01, GENASEX, from the French National Research Agency.

## Appendix 1

### MinION assembly

We first produced a *de novo* assembly of MinION reads with wtdbg2 (Ruan and Li 2020). Reads that were not used were merged with the obtained contigs. 10X Illumina reads were mapped to this MinION assembly with bwa mem. The resulting bam files were used to correct the MinION assembly with Pilon (Walker et al. 2014).

### 10X assembly

We then used 10X Illumina reads to produce a second *de novo* assembly with Supernova (Weisenfeld et al. 2018). Two pseudohaplotypes were created with the “mkoutput” command. 10X reads were then mapped against the first pseudohaplotype with bowtie2 (Langmead et al. 2009). The resulting bam file was used to remove duplicated contigs and cut overlapping parts of the supernova pseudohaplotype with “Purge_haplotigs”. Finally, we used Links (Warren et al. 2015) to link the contigs of this purged assembly with the MinION corrected contigs to get our final reference assembly (Tab S1).

**Table S1.** Characteristics of the assembly. N represents the number of scaffolds included in the calculation.

| Measure | Result | N |
| --- | --- | --- |
| Sum | 1247144234 | 129396 |
| N50 | 27463 | 11676 |
| N60 | 19082 | 17114 |
| N70 | 12065 | 25372 |
| N80 | 7247 | 38763 |
| N90 | 3605 | 62956 |
| N100 | 102 | 129396 |

### BUSCO

We used BUSCO (Manni et al. 2021) to check for completeness by comparing to the arthropod database. This analysis is summarized in Tab S2.

**Table S2.** Results of the BUSCO analysis.

| BUSCOs | Result | Percentage |
| --- | --- | --- |
| Complete BUSCOs | 491 | 48.5% |
| Complete and single-copy BUSCOs | 476 | 47.0% |
| Complete and duplicated BUSCOs | 15 | 1.5% |
| Fragmented BUSCOs | 156 | 15.4% |
| Missing BUSCOs | 366 | 36.1% |
| Total BUSCO groups searched | 1013 |  |

## Appendix 2.

### Autosomal LOH analysis

We analysed the LOH data using likelihood. We note *q* the proportion of autosomal scaffolds losing heterozygosity. We assume that *q* values are Beta distributed with parameters *a* and *b*, and that these parameters can vary between males and females. We note *n*_*i*_ and *m*_*i*_ the observed number of scaffolds that loose or retain heterozygosity for individual *i*. We note **n** and **m** the vector of all *n*_*i*_ and *m*_i_ and □ the vector of parameters to be estimated. The likelihood of the data can then be written

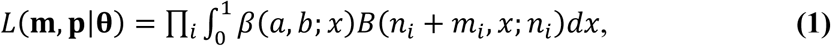

where □ (*a*□ □ *b* ; *x*) denotes the probability to draw *x* in a Beta distribution with parameters *a* and *b* and where □ (*n, x*; *k*) denotes the probability to draw *k* success among *n* trials with a probability of success *x* (i.e. in a binomial distribution with parameters *n* and *x*). We estimated a and b parameters independently for males and females and, in another model where a and b were constrained to be identical between males and females. We tested whether male and female LOH differed by comparing these models with a likelihood ratio test. These tests were done using Mathematica v.9 (Wolfram Research 2012).

## REFERENCES

Abatzopoulos, T. J. 2018. The repeated emergence of asexuality, the hidden genomes and the role of parthenogenetic rare males in the brine shrimp Artemia. J. Biol. Res. 1–5. BioMed Central.

Abreu-Grobois, F., and J. Beardmore. 2001. The generation of males by diploid parthenogenetic Artemia cannot occur in the way Stefani suggested. P. 1 in A. Maeda-Martinez, B. Timms, D. Rogers, G. Murugan, and A. Abreu-Grobois, eds. Proceedings of the 4th International Large Branchiopod Symposium. La Paz, Baja California, Mexico.

Accioly, I. V., I. M. C. Cunha, J. C. M. Tavares, and W. F. Molina. 2014. Chromosome Banding in Crustacea. I. Karyotype, Ag-NORs, C Banding and Treatment with EcoRI, PstI and KpnI Restriction Endonucleases in Artemia franciscana. Biota Amaz. 4:15–19.

Archetti, M. 2010. Complementation, genetic conflict, and the evolution of sex and recombination. J. Hered. 101:1–13.

Archetti, M. 2004. Recombination and loss of complementation: A more than two-fold cost for parthenogenesis. J. Evol. Biol. 17:1084–1097.

Baird, N. A., P. D. Etter, T. S. Atwood, M. C. Currey, A. L. Shiver, Z. A. Lewis, E. U. Selker, W. A. Cresko, and E. A. Johnson. 2008. Rapid SNP discovery and genetic mapping using sequenced RAD markers. PLoS One 3:1–7.

Barigozzi, C. 1974. Artemia: a survey of its significance in genetic proplems. P. in T. Dobzhansky, M. Hecht, and W. Steere, eds. Evolutionary Biology. Springer, Boston, MA.

Baxevanis, A. D., I. Kappas, and T. J. Abatzopoulos. 2006. Molecular phylogenetics and asexuality in the brine shrimp Artemia. Mol. Phylogenet. Evol. 40:724–738.

Blackman, R. L. 1972. The inheritance of life-cycle differences in Myzus persicae (Sulz.) (Hem., Aphididae). Bull. Entomol. Res. 62:281–294.

Blackman, R. L., and D. F. Hales. 1986. Behaviour of the X chromosomes during growth and maturation of parthenogenetic eggs of Amphorophora tuberculata (Homoptera, Aphididae), in relation to sex determination. Chromosoma 94:59–64.

Bolger, A. M., M. Lohse, and B. Usadel. 2014. Trimmomatic: A flexible trimmer for Illumina sequence data. Bioinformatics 30:2114–2120.

Bowen, S. T. 1963. The Genetics of Artemia Salina. II. White Eye, a Sex-Linked Mutation. Biol. Bull. 124:17–23.

Boyer, L., J.-Z. R.M. Mosna, C. R. Haag, T. Lenormand, R. Zahab, M. Mosna, C. R. Haag, and T. Lenormand. 2021. Not So Clonal Asexuals: Unraveling The Secret Sex Life Of Artemia parthenogenetica. Evol. Lett. 5:164–174.

Broman, K. W., H. Wu, S. Sen, and G. A. Churchill. 2003. R/qtl: QTL mapping in experimental crosses. Bioinformatics 19:889–890.

Browne, R. A., and C. W. Hoopes. 1990. Genotype Diversity and Selection in Asexual Brine Shrimp (Artemia). Evolution 44:1035–1051.

Card, D. C., F. J. Vonk, S. Smalbrugge, N. R. Casewell, W. Wüster, T. A. Castoe, G. W. Schuett, and W. Booth. 2021. Genome-wide data implicate terminal fusion automixis in king cobra facultative parthenogenesis. Sci. Rep. 11:1–9.

Catchen, J., P. A. Hohenlohe, S. Bassham, A. Amores, and W. A. Cresko. 2013. Stacks: An analysis tool set for population genomics. Mol. Ecol. 22:3124–3140.

Danecek, P., A. Auton, G. Abecasis, C. A. Albers, E. Banks, M. A. DePristo, R. E. Handsaker, G. Lunter, G. T. Marth, S. T. Sherry, G. McVean, and R. Durbin. 2011. The variant call format and VCFtools. Bioinformatics 27:2156–2158.

de Vos, S., P. Bossier, G. van Stappen, I. Vercauteren, P. Sorgeloos, and M. Vuylsteke. 2013. A first AFLP-Based Genetic Linkage Map for Brine Shrimp Artemia franciscana and Its Application in Mapping the Sex Locus. PLoS One 8:1–10.

Dobin, A., and T. R. Gingeras. 2015. Mapping RNA-seq reads with STAR. Curr. Protoc. Bioinform. 51.

Doums, C., C. Ruel, J. Clémencet, P. Fédérici, L. Cournault, and S. Aron. 2013. Fertile diploid males in the ant Cataglyphis cursor: A potential cost of thelytoky? Behav. Ecol. Sociobiol. 67:1983–1993.

Emms, D. M., and S. Kelly. 2015. OrthoFinder: solving fundamental biases in whole genome comparisons dramatically improves orthogroup inference accuracy. Genome Biol. 16:1–14. Genome Biology.

Engelstädter, J. 2017. Asexual but Not Clonal : Evolutionary Processes in Automictic Populations. Genetics 206:993–1009.

Engelstädter, J. 2008. Constraints on the evolution of asexual reproduction. BioEssays 30:1138–1150.

Engelstädter, J., C. Sandrock, and C. Vorburger. 2011. Contagious parthenogenesis, automixis, and a sex determination meltdown. Evolution 65:501–511.

Goudie, F., M. H. Allsopp, M. Beekman, P. R. Oxley, J. Lim, and B. P. Oldroyd. 2012. Maintenance and loss of heterozygosity in a thelytokous lineage of honey bees (Apis mellifera capensis). Evolution 66:1897–1906.

Haag, C. C. R., L. Theodosiou, R. Jabbour-Zahab, T. Lenormand, R. Zahab, and T. Lenormand. 2017. Low recombination rates in sexual species and sex-asex transitions. Philos. Trans. R. Soc. London 372:20160461.

Hebert, P. D. N., and T. Crease. 1983. Clonal diversity in populations of Daphnia Pulex reproducing by obligate parthenogenesis. Heredity 51:353–369.

Huylmans, A. K., M. A. Toups, A. MacOn, W. J. Gammerdinger, and B. Vicoso. 2019. Sex-biased gene expression and dosage compensation on the artemia franciscana Z-chromosome. Genome Biol. Evol. 11:1033–1044.

Innes, D. J., C. J. Fox, and G. L. Winsor. 2000. Avoiding the cost of males in obligately asexual Daphnia pulex (Leydig). Proc. R. Soc. B Biol. Sci. 267:991–997.

Innes, D. J., and P. D. N. Hebert. 1988. The Origin and Genetic Basis of Obligate Parthenogenesis in Daphnia pulex. Evolution 42:1024–1035.

Kim, D., J. M. Paggi, C. Park, C. Bennett, and S. L. Salzberg. 2019. Graph-based genome alignment and genotyping with HISAT2 and HISAT-genotype. Nat. Biotechnol. 37:907– 915.

Knaus, B. J., and N.J. Grünwald. 2017. vcfr: a package to manipulate and visualize variant call format data in R. Mol. Ecol. Resour. 17:44–53.

Korneliussen, T. S., A. Albrechtsen, and R. Nielsen. 2014. ANGSD: Analysis of Next Generation Sequencing Data. BMC Bioinformatics 15:1–13.

Lenormand, T., J. Engelstädter, S. E. Johnston, E. Wijnker, and C. R. Haag. 2016. Evolutionary mysteries in meiosis. Philos. Trans. R. Soc. B Biol. Sci. 371.

Li, H., and R. Durbin. 2009. Fast and accurate short read alignment with Burrows-Wheeler transform. Bioinformatics 25:1754–1760.

Li, H., B. Handsaker, A. Wysoker, T. Fennell, J. Ruan, N. Homer, G. Marth, G. Abecasis, and R. Durbin. 2009. The Sequence Alignment/Map format and SAMtools. Bioinformatics 25:2078–2079.

Lynch, M. 1984. Destabilizing hybridization, general-purpose genotypes and geographic parthenogenesis. Q. Rev. Biol. 59:257–290.

Maccari, M., F. Amat, F. Hontoria, and A. Gómez. 2014. Laboratory generation of new parthenogenetic lineages supports contagious parthenogenesis in Artemia. PeerJ 2:e439.

Maccari, M., A. Gómez, F. Hontoria, and F. Amat. 2013. Functional rare males in diploid parthenogenetic Artemia. J. Evol. Biol. 26:1934–1948.

MacDonald, G. H., and R. A. Browne. 1987. Inheritance and reproductive role of rare males in a parthenogenetic population of the brine shrimp, Artemia parthenogenetica. Genetica 75:47–53.

Maruki, T., and M. Lynch. 2015. Genotype-frequency estimation from high-throughput sequencing data. Genetics 201:473–486.

Neiman, M., K. Larkin, A. R. Thompson, and P. Wilton. 2012. Male offspring production by asexual Potamopyrgus antipodarum, a New Zealand snail. Heredity 109:57–62.

Nougué, O., N. O. Rode, R. Jabbour-zahab, A. Ségard, L. M. Chevin, C. R. Haag, and T. Lenormand. 2015. Automixis in Artemia: Solving a century-old controversy. J. Evol. Biol. 28:2337–2348.

Olsen, M., and S. Marsden. 1954. Natural parthenogenesis in turkey eggs. Science 120:545– 546.

Paris, J. R., J. R. Stevens, and J. M. Catchen. 2017. Lost in parameter space: a road map for stacks. Methods Ecol. Evol. 8:1360–1373.

Parraguez, M., G. Gajardo, and J. A. Beardmore. 2009. The new world artemia species A. franciscana and A. persimilis are highly differentiated for chromosome size and heterochromatin content. Hereditas 146:93–103.

Pijnacker, L. P., and M. A. Ferwerda. 1980. Sex chromosomes and origin of males and sex mosaics of the parthenogenetic stick insect Carausius morosus Br. Chromosoma 79:105– 114.

Ranwez, V., S. Harispe, F. Delsuc, and E. J. P. Douzery. 2011. MACSE: Multiple alignment of coding SEquences accounting for frameshifts and stop codons. PLoS One 6.

Rastas, P. 2017. Lep-MAP3: Robust linkage mapping even for low-coverage whole genome sequencing data. Bioinformatics 33:3726–3732.

Rispe, C., J. Bonhomme, and J. C. Simon. 1999. Extreme life-cycle and sex ratio variation among sexually produced clones of the aphid Rhopalosiphum padi (Homoptera: Aphididae). Oikos 254–264.

Rochette, N. C., and J. M. Catchen. 2017. Deriving genotypes from RAD-seq short-read data using Stacks. Nat. Protoc. 12:2640–2659. Nature Publishing Group.

Rochette, N. C., A.G. Rivera-Colón, and J. M. Catchen. 2019. Stacks 2: Analytical methods for paired-end sequencing improve RADseq-based population genomics. Mol. Ecol. 28:4737–4754.

Rode, N. O., R. Jabbour-Zahab, L. Boyer, É. Flaven, F. Hontoria, G. Van Stappen, F. Dufresne, C. Haag, and T. Lenormand. in press. The origin of asexual brine shrimps. Am. Nat.

Sandrock, C., and C. Vorburger. 2011. Single-Locus recessive inheritance of asexual reproduction in a parasitoid wasp. Curr. Biol. 21:433–437.

Simon, J. C., F. Delmotte, C. Rispe, and T. Crease. 2003. Phylogenetic relationships between parthenogens and their sexual relatives: The possible routes to parthenogenesis in animals. Biol. J. Linn. Soc. 79:151–163.

Simon, J. C., N. Leterme, and A. Latorre. 1999. Molecular markers linked to breeding system differences in segregating and natural populations of the cereal aphid Rhopalosiphum padi L. Mol. Ecol. 8:965–973.

Smit, A., R. Hubley, and P. Green. 2013. RepeatMasker. WA: Institute for Systems Biolog, Seattle.

Smith, R. J., T. Kamiya, and D. J. Horne. 2006. Living males of the “ancient asexual” Darwinulidae (Ostracoda: Crustacea). Proc. R. Soc. B Biol. Sci. 273:1569–1578.

Snyder, D. W., C. H. Opperman, and D. McK. Bird. 2006. A Method for Generating Meloidogyne incognita Males. J. Nematol. 38:192–194.

Stanke, M., O. Schöffmann, B. Morgenstern, and S. Waack. 2006. Gene prediction in eukaryotes with a generalized hidden Markov model that uses hints from external sources. BMC Bioinformatics 7:1–11.

Stefani, R. 1963. Il centromero non localizzato in Artemia salina Leach. Rend. Accad. Naz. Lincei Cl. Sci. fis. mat. nat 35:375–378.

Stefani, R. 1964. The origin of males in parthenogenetic populations of Artemia salina. Riv. Biol. 57:147.

Strunnikov, V. 1995. Control over reproduction, sex, and heterosis of the silkworm. Harwood Academic Publishers., Luxembourg.

Svendsen, N., C. M. O. Reisser, M. Dukić, V. Thuillier, A. Ségard, C. Liautard-Haag, D. Fasel, E. Hürlimann, T. Lenormand, Y. Galimov, and C. R. Haag. 2015. Uncovering cryptic asexuality in Daphnia magna by RAD-sequencing. Genetics 201:1143–1155. Genetics.

R Core Team 2021. R: A language and environment for statistical computing. R Foundation for Statistical Computing, Vienna, Austria.

van der Auwera, G. A., and B.D. O’Connor. 2020. Genomics in the Cloud: Using Docker, GATK, and WDL in Terra. 1st ed.

van der Kooi, C. J., and T. Schwander. 2014a. Evolution of asexuality via different mechanisms in grass thrips (thysanoptera: Aptinothrips). Evolution 68:1883–1893.

van der Kooi, C. J., and T. Schwander. 2014b. On the fate of sexual traits under asexuality. Biol. Rev. 89:805–819. John Wiley & Sons, Ltd (10.1111).

Watts, P. C., K. R. Buley, S. Sanderson, W. Boardman, C. Ciofi, and R. Gibson. 2006. Parthenogenesis in Komodo dragons. Nature 444:1021–1022.

Wilson, A. C. C., P. Sunnucks, and D. F. Hales. 1997. Random loss of X chromosome at male determination in an aphid, Sitobion near fragariae, detected using an X-linked polymorphic microsatellite marker. Genet. Res. 69:233–236.

Yang, Z. 1997. Paml: A program package for phylogenetic analysis by maximum likelihood. Bioinformatics 13:555–556.

## APPENDIX REFERENCES

Langmead, B., C. Trapnell, M. Pop, and S. L. Salzberg. 2009. Ultrafast and memory-efficient alignment of short DNA sequences to the human genome. Genome Biol. 10.

Manni, M., M. R. Berkeley, M. Seppey, F. A. Simão, and E. M. Zdobnov. 2021. BUSCO Update: Novel and Streamlined Workflows along with Broader and Deeper Phylogenetic Coverage for Scoring of Eukaryotic, Prokaryotic, and Viral Genomes. Mol. Biol. Evol. 38:4647–4654.

Ruan, J., and H. Li. 2020. Fast and accurate long-read assembly with wtdbg2. Nat. Methods 17:155–158.

Walker, B. J., T. Abeel, T. Shea, M. Priest, A. Abouelliel, S. Sakthikumar, C. A. Cuomo, Q. Zeng, J. Wortman, S. K. Young, and A. M. Earl. 2014. Pilon: An integrated tool for comprehensive microbial variant detection and genome assembly improvement. PLoS One 9.

Warren, R. L., C. Yang, B. P. Vandervalk, B. Behsaz, A. Lagman, S. J. M. Jones, and I. Birol. 2015. LINKS: Scalable, alignment-free scaffolding of draft genomes with long reads. Gigascience 4. GigaScience.

Weisenfeld, N. I., V. Kumar, P. Shah, D. M. Church, and D. B. Jaffe. 2018. Direct determination of diploid genome sequences. Genome Res. 28:757–767.

Wolfram Research, I. 2012. Mathematica 9.0. Wolfram Research, Inc., Champaign, IL.

