## Supplementary figures for "Asexual male production by ZW recombination in *Artemia parthenogenetica*"

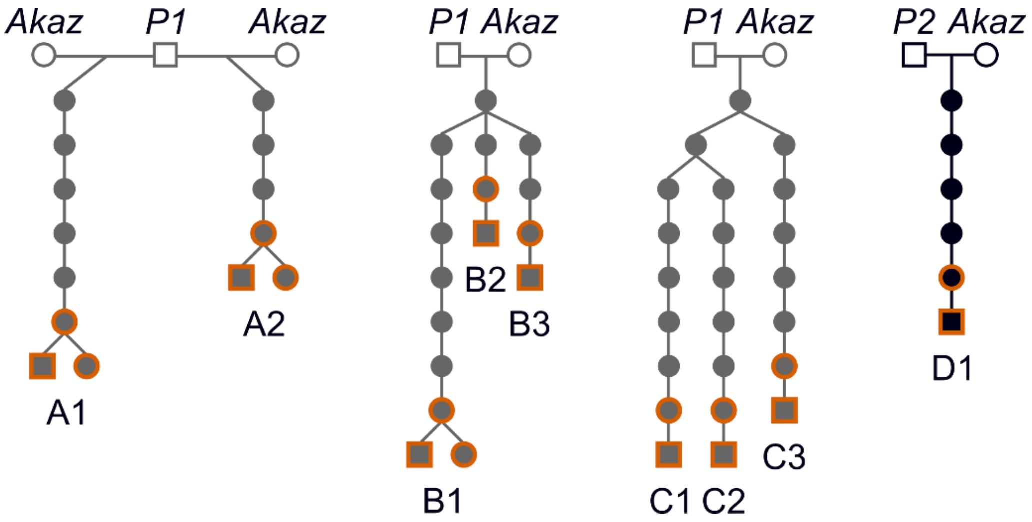


**Figure S1. Origin of the individuals studied.** Circles represent female individuals and squares represent male individuals. Open circles and squares represent the sexual females and rare males used to generate new asexual lineages. Lineages in grey were produced by rare males originating from the P1 (Aigues-Mortes) population, while the lineage in black was produced by a rare male from the P2 (Lake Urmia) population. The individuals included in the study are highlighted in orange.

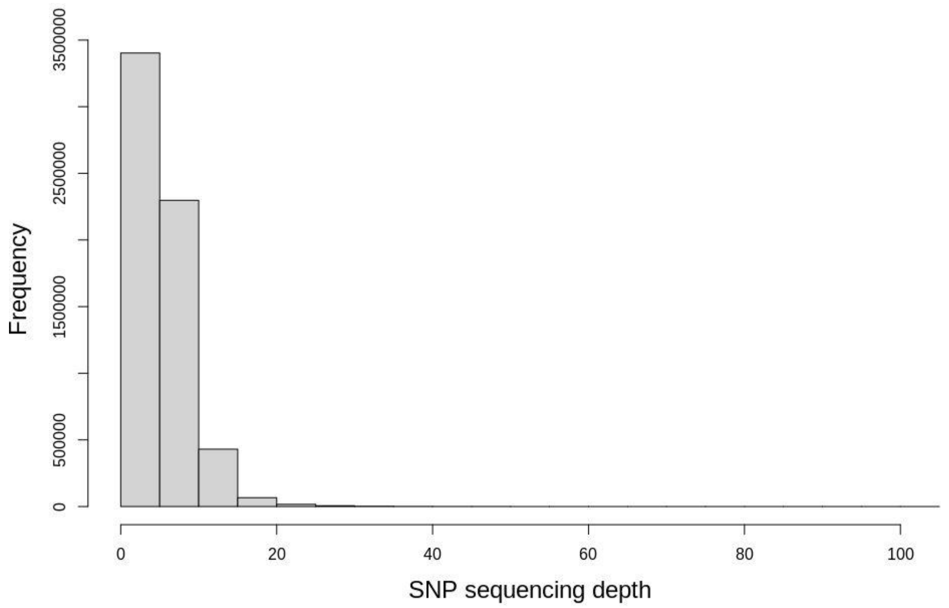


**Figure S2. Distribution of sequencing depth per SNP.**

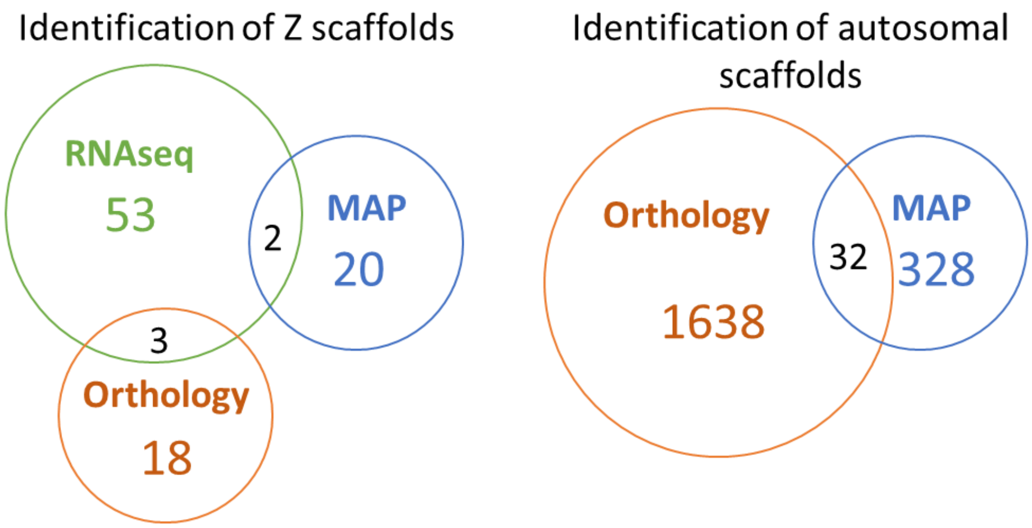


**Figure S3. Identification of Z and autosomal scaffolds by the different methods.** RNAseq refers to the comparison of SNP heterozygosity between males and females. MAP refers to the assignation of scaffolds to the linkage map of *Akaz*. Orthology refers to the assignation of scaffolds based on the autosomal/Z assignation made by Huylmans et al. 2019.

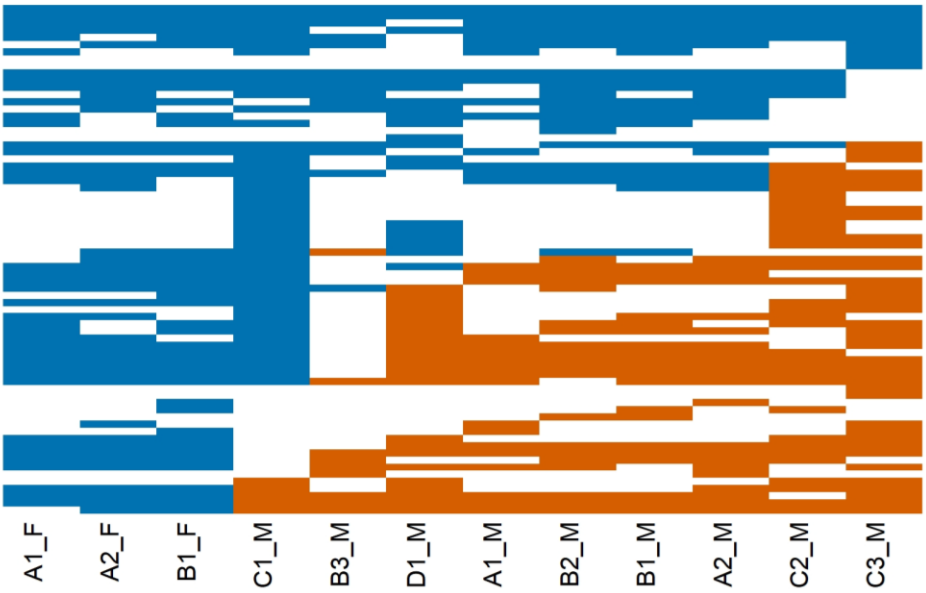


**Figure S4. LOH on the sex chromosome scaffolds after removal of scaffolds identified by orthology with *A. franciscana*.** Figure construction is the same as Figure 2 (main text). The removal of the 6 scaffolds did not affect the pattern observed on the figure.

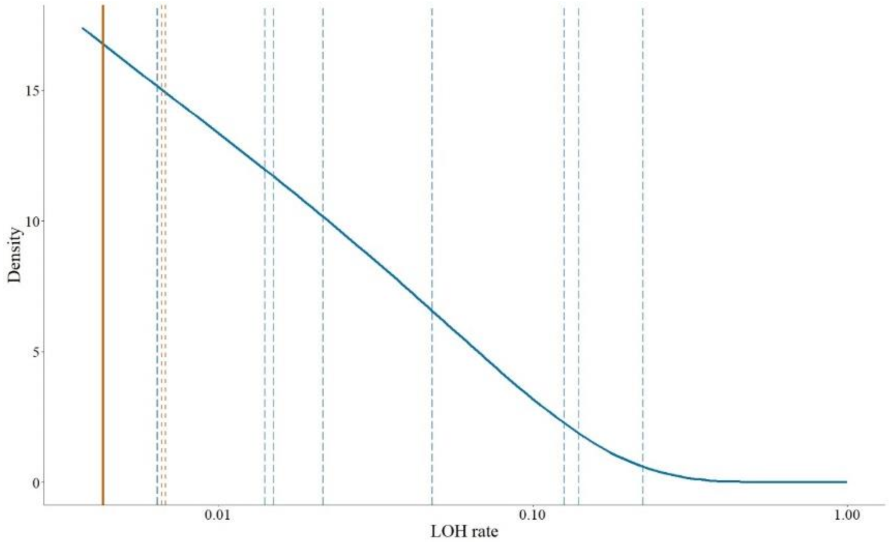


**Figure S5. LOH rate on 260 autosomal scaffolds in male (blue) and female (orange) offspring after removal of scaffolds identified by orthology with *A. franciscana*.** Dashed lines represent LOH rates in autosomal scaffolds for individual offspring, and solid lines represent the estimated distributions from our best model. One of the females presented an LOH rate of 0, and is thus not represented on this log-scaled figure. The scaffold removal largely did not affect the results. The best model of autosomal LOH distribution among offspring also displayed a difference between males and females (likelihood ratio test, $P = 0.03$). In males, LOH rate followed a beta distribution of parameters a=0.80 and b=11.2. In females, LOH rate followed a beta distribution with an estimated mean of $4.10^{-3}$ and a near-zero variance of $10^{-9}$.
